# Predicting genotypic values associated with gene interactions using neural networks: A simulation study for investigating factors affecting prediction accuracy

**DOI:** 10.1101/2019.12.18.881912

**Authors:** Akio Onogi

## Abstract

Genomic prediction has been applied to various species of plants and livestock to enhance breeding efficacy. Neural networks including deep neural networks are attractive candidates to predict phenotypic values. However, the properties of neural networks in predicting non-additive effects have not been clarified. In this simulation study, factors affecting the prediction of genetic values associated with gene interactions (*i.e.*, epistasis) were investigated using multilayer perceptron. The results suggested that (1) redundant markers should be pruned, although markers in LD with QTLs are less harmful, (2) predicting epistatic genetic values with neural networks in real populations would be infeasible using training populations of 1000 samples, (3) neural networks with two or fewer hidden layers and a sufficient number of units per hidden layer would be useful, particularly when a certain number of interactions is involved, and (4) neural networks have greater capability to predict epistatic genetic values than random forests, although neural networks are more sensitive to training population size and the level of epistatic genetic variance. These lessons also would be applicable to other regression problems in which interactions between explanatory variables are expected, *e.g.*, gene-by-environment interactions.

## Introduction

Genomic prediction/selection [1] has been applied to plants and livestock to enhance breeding efficacy [2–4]. Genomic prediction is also used in medical fields to predict the risk scores of polygenic diseases [5]. In genomic prediction applied to breeding, the purpose is often to predict the additive components of genotypic values *i.e.*, breeding values, of new varieties or animals because the breeding value is the value transmitted from parents to offspring. In addition, this is also because non-additive (*i.e.*, dominance and epistatic) genetic variances are often small in real populations because these variances decrease as the minor allele frequencies of quantitative trait loci (QTLs) approach zero [6]. Nevertheless, predicting non-additive components of genotypic values remains of interest in breeding because the minor allele frequencies of QTLs can be high in synthetic populations and because obtaining favorable phenotypic values is the goal in production at farms.

Neural networks have been applied for genomic prediction as attractive candidates to capture non-additive genetic values because of their capability to approximate complex functions [7–13]. For example, neural networks with a single hidden layer of 1–6 units (neurons) were used to predict the phenotypic values (individual outcomes) of wheat and dairy cows and it was found that the prediction accuracy was higher than that of linear approaches [8]. Whereas the authors used the additive genetic relationship matrices as inputs, another study used SNP genotypes as inputs and reported that single hidden layer neural networks with 1–4 units displayed nearly equivalent accuracy as Bayesian regression methods in predicting the expected progeny differences of marbling scores in beef cattle [11]. In the prediction of days to heading and grain yield of wheat, radial basis function neural networks and the reproducing kernel Hilbert space regression (RKHS) generally exhibited greater predictive accuracy than linear regression methods [12]. Genomic prediction using relatively small neural networks is reviewed in ref. [14].

Because convolutional neural networks displayed remarkable performance in ImageNet large scale visual recognition challenge 2012 [15,16], deep neural networks have attracted tremendous attention in various application fields such as vision object recognition and natural language processing [17,18], and they have also been applied to genomic prediction. Convolutional neural networks were used to predict the agronomic traits of wheat [19] and quantitative traits of humans including height [20]. Multilayer perceptron was applied to the agronomic traits of wheat, maize, and *Arabidopsis* data [21] and human quantitative traits [20]. The predictive accuracies of the deep neural networks reported in these studies were generally comparable to those of linear regression methods, and these networks failed to display the remarkable performance reported in other applications such as image recognition.

The real population data used in the aforementioned studies differed regarding various factors including the level of non-additive genetic variances, the number of QTLs, and linkage disequilibrium (LD) between QTLs and markers. Moreover, in real data analyses, it is impossible to enlarge the training population size beyond the available size. Thus, it is difficult to investigate the influences of these factors on the capability of neural networks to predict non-additive genetic values from only real data analyses. In other words, even if neural networks fail to show better performance than other methods, the causes are unclear in real data analyses. In this study, I examined the influences of these factors on the predictive performance of neural networks using simulations and attempted to clarify the suitable network structures for predicting non-additive genetic values.

Compared with dominance genetic values, predicting epistatic values is expected to be difficult because the number of combinations of QTLs/markers to be searched exponentially increases as the numbers of QTLs/markers increase. Consequently, the neural networks required for epistatic values are expected to be more complex or larger than those for dominance values. Moreover, predicting epistatic values can be regarded as a more general task than that of dominance values because interactions of explanatory variables in regression problems can be found in various application fields. Thus, in this study, considering the computational costs and available computational resources, I focused on predicting epistatic genetic effects.

## Materials and Methods

### Simulation scenarios

Because intensive simulations using deep neural networks are time-consuming, F_2_ populations of selfing crops (e.g., rice and soybean) for QTL mapping where relatively small numbers of DNA markers (two to three hundred) are often genotyped [22,23] were assumed in this study. The number of segregated markers was assumed to be 200. Because of F_2_ populations, the allele frequencies of QTLs were 0.5, which maximizes the epistatic variance.

The additive-by-additive model without dominance was assumed [6]. Among the 200 markers, 20 were selected as QTLs or were assumed to be tightly linked with QTLs, and 10 interacting pairs were generated. In the scenario of Q20UM180, the remaining 180 markers were not in LD with any QTL. But in scenario Q20LM180, the 180 markers were in LD with QTLs. Each QTL was accompanied by nine linked markers, resulting in 20 blocks, comprising one QTL and nine linked markers. To make the comparison between scenarios more meaningful, two unrealistic scenarios were also assumed. In the first scenario (Q20), only genotypes of 20 QTLs were used for prediction. In the second scenario (Q200), the 200 markers were all assumed to be QTLs or tightly linked to QTLs, and comprised 100 interacting pairs. These scenarios were used to investigate the influence of redundant markers and epistatic complexity (i.e., numbers of interacting pairs of QTLs) on prediction accuracy. As demonstrated later, comparisons of the four scenarios Q20UM180, Q20LM180, Q20, and Q200 revealed properties of neural networks on predicting epistatic genetic values and indicated the feasibility of the method for predicting real data.

Large training population sizes (1000, 5000, and 10,000) were assumed so that the performances of neural networks could be compared, although the sizes of F_2_ populations for QTL mapping were usually several hundreds. Three broad-sense heritabilities (1.0, 0.6, and 0.2) were also assumed. Thus, in total, 36 (4 QTL-marker scenarios × 3 training population sizes × 3 broad-sense heritabilities) scenarios were considered. A testing population (*n* = 1000) was also simulated for each scenario to aid learning of the multilayer perceptron. Each scenario was replicated three times. Fluctuations of prediction accuracies between replicates were small in most simulation scenarios, as shown in the Results and Discussion section.

### Genotype simulation

The genotypes of QTLs and markers in Q20UM180, Q20, and Q200 were simulated using a Bernoulli trial. The allele frequency was 0.5 for all QTLs and markers. The genotypes in Q20LM180 were simulated using a multivariate normal distribution. For each block consisting of one QTL and nine markers, random vectors were generated from a multivariate normal distribution of which the mean was a 0-vector of length 10, and the (co)variance was a 10 × 10 symmetric matrix with diagonals of 1 and off-diagonals of 0.7. The number of random vectors was the summation of the training and testing population sizes. Then, for each QTL or marker, values in the first quantile were regarded as homozygous (0), those in the second and third quantiles were considered heterozygous (1), and the remaining were deemed homozygous of another allele (2), assuming that the allele frequency was 0.5. On average, *r*^*2*^ between the QTL and markers within a block was 0.34 ± 0.02.

### Genotypic and phenotypic values

The effect of the additive-by-additive epistasis was generated from the standard normal distribution. Because additive effects were not simulated, genotypic values were generated only via epistatic effects. Residual errors were generated from a normal distribution of which the mean was 0 and the variance was calculated using the broad-sense heritability and empirical variance of genotypic values. Hereafter, I refer to the broad-sense heritability as heritability for the sake of simplification. Phenotypic values were generated by summing genotypic values and residual errors.

### Neural networks

Multilayer perceptron implemented in Tensorflow (ver. 1.12.0) was used from the Keras library (ver. 2.2.4). I used desktop machines equipped with two GPU nodes (GeForce GTX 1080 Ti, Nvidia, CA, USA) per machine. The operating system was Ubuntu 16.04 Lite, and the scripts were written with python ver. 3.5.2. The number of units in the input layer was the number of covariates, *i.e.*, the number of QTLs and markers (20 or 200). The genotypes of QTLs and markers were coded additively as 0 (aa), 1 (Aa), or 2 (AA). The number of hidden layers was 1, 2, 3, or 4, and the number of units per hidden layer was 20, 200, 400, 800, or 1200, resulting 20 (4 × 5) models. The activation function in the hidden layers was the rectified linear unit [24], and the optimizer was Adam [25]. The learning rates of Adam were the default values. The loss function was the mean squared error (MSE). The initial values of weights were generated using the Glorot uniform initializer [26], which is the default choice of the function Dense of Keras. The initial value of bias terms was 0. The batch normalization technique [27] was implemented for each layer. Dropout [28] was tested for simulation scenarios with a heritability of 0.6 for Q20UM180 and models with two hidden layers. The rates were 0.1, 0.2, and 0.3. One-hot encoding of genotypes was also tested for the simulation scenarios with a heritability of 0.6 for Q20UM180 and models with two hidden layers. In this coding, three covariates were made for each QTL/marker, and each covariate represents the presence (1) or absence (0) of genotypes aa, Aa, and AA. Thus, the number of units in the input layer was 600.

The number of epochs was set to 1000, and the batch size was 32. While training at each epoch, 20% of the training population was randomly chosen and used for validation. The training results (*i.e.*, weights of neural networks) were stored for each 10 epochs, and the result with the smallest MSE between predictions and phenotypic values in the validating population was used for prediction. Phenotypic values were standardized when training.

### Model evaluation

Models were evaluated by comparing the predicted values with the true genotypic values in the testing population. Because the correlation between the predicted and true values is related to the selection response in the breeder’s equation, I report the Pearson’s product moment correlation coefficient as the metric to assess the predictive accuracy in this study.

As an alternative method to predict epistatic genotypic values, I also evaluated random forests [29] implemented in an R package ranger [30]. The number of trees (*num.trees*) was 10,000. The number of variables used for constructing a tree (*mtry*) was 4, 14, 20, 40, or 60 for Q20UM180, Q20UL180, and Q200 and 2, 4, 8, or 16 for Q20. The best *mtry* values were chosen via internal 5-fold cross-validation in the training population. Although RKHS is another popular method for predicting non-additive effects, considering the computational burden for hyperparameter (*e.g.*, kernel types and bandwidth) tuning using cross-validation, particularly in *n* = 10,000 scenarios, I avoided comparisons with RKHS.

## Results and Discussion

### Influences of the genetic architecture, LD, and training population size

The numbers of QTLs and markers as well as LD between them affected the correlation coefficients, *i.e.*, prediction accuracies (Figures 1–3). Generally, the accuracies were highest for the Q20 scenario. Although adding markers that were in LD with QTLs decreased the accuracies (Q20LM180 scenario), the markers that were not in LD with QTLs affected the accuracies with greater magnitudes (Q20UM180 scenario), particularly when the training population size was small (1000). The undesirable influence of unlinked markers on prediction using neural networks was also reported by a previous study [10]. Thus, it is highly recommended to prune markers before feeding them to neural networks. LD between markers and QTLs can mitigate the influence of additional markers in the prediction of epistatic genetic values, which was demonstrated for the first time in the present study. This is probably because these markers would be helpful in searching interactions among QTLs. The accuracies were generally lowest for the Q200 scenario. This result is reasonable considering the increased number of interactions.

**Figure 1.**
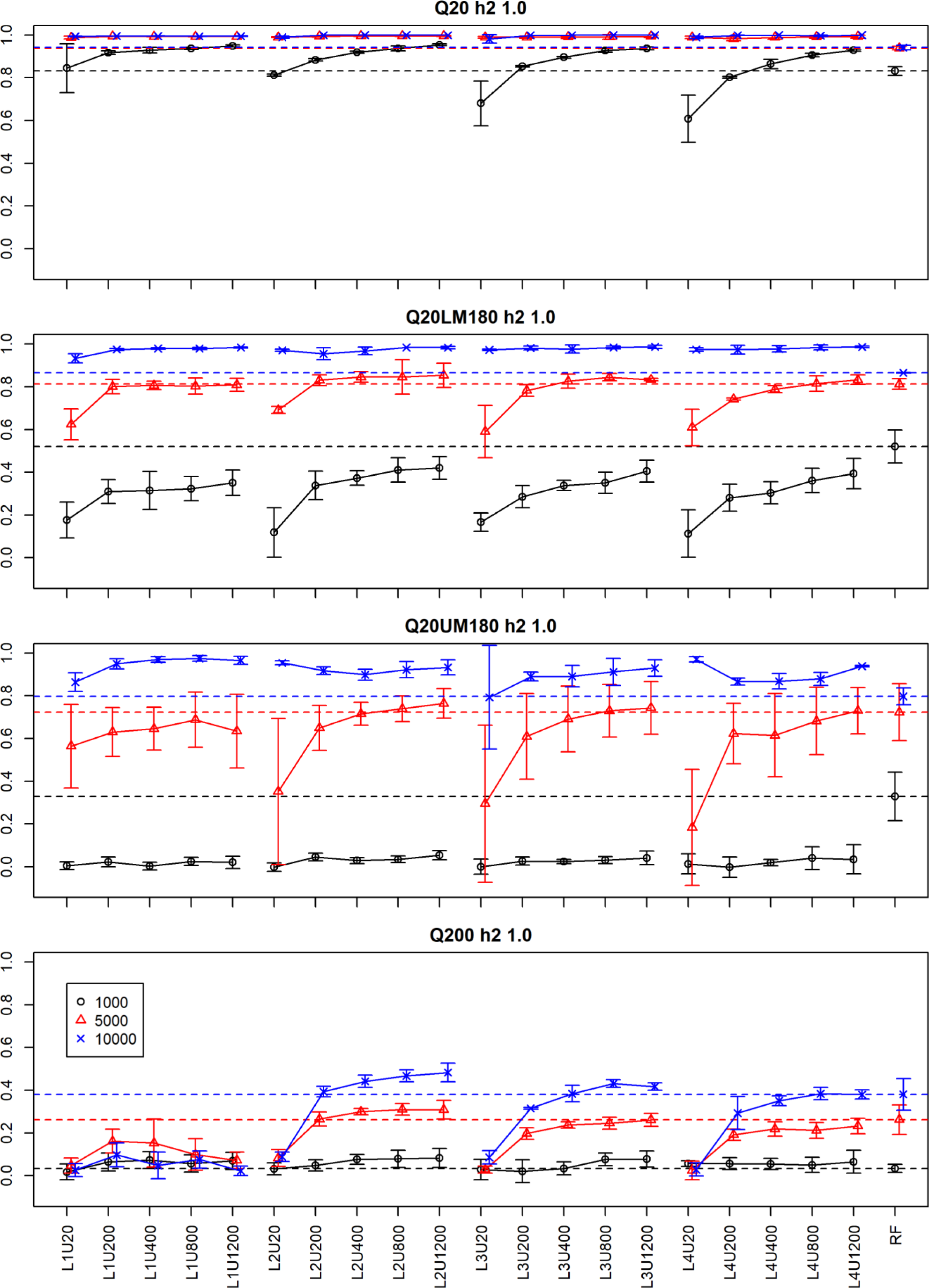
Prediction accuracies when the broad-sense heritability was 1.0. The four plots correspond with the four quantitative trait locus (QTL)-marker scenarios: Q20, Q20LM180, Q20UM180, and Q200 (see the text for the explanations of these QTL-marker scenarios). The y-axis is the prediction accuracy, and the x-axis presents the neural network models (L1U20 to L4U1200) and random forests (RFs). L and U in the model names represent the number of hidden layers and units per hidden layer, respectively. The dashed horizontal lines indicate the mean accuracies obtained using RFs. Colors and symbols indicate the training population sizes (1000, 5000, and 10,000). Bars are the standard deviations.

Heritability and training population size also affected the prediction accuracy, as the accuracies tended to decrease as these values decreased. These tendencies are generally observed in genomic prediction regardless of the methods used [31–33]. With a training population size of 1000, neural networks were unable to display even moderate accuracy (~0.5) except for the Q20 scenario, which is the most unrealistic (or ideal) scenario (Figures 1–3). Training population sizes are often less than 1000 in plant studies of genomic prediction [34,35]. Considering the greater numbers of QTLs and markers included in real populations, our results suggest that predicting epistatic genetic values using neural networks is often infeasible for plant data sets, although predicting non-additive effects is useful to select superior genotypes practically in clonally propagated plants.

### Influences of the model structures

Increasing the number of hidden layers from one to two was effective when the training population size was 5000 or 10,000 in Q200 (Figures 1–3), although further increasing the layers to three or four did not increase the accuracy. These results suggest that multiple hidden layers are effective as the number of interacting QTLs increases; however, it cannot be concluded that two hidden layers are sufficient even when the number of interacting QTLs increases more (*e.g.*, 1000). The results also suggest that deeper neural networks require greater training population sizes, which is reasonable considering the growing number of parameters. In each scenario, increasing the number of units did not result in higher prediction accuracies when high accuracies (>0.8) were already achieved with the fewest number of units (20 in this study). However, when this was not the case, the accuracy tended to increase as the number of units increased. In a previous study, three multilayer perceptrons designed for additive genotype coding were compared using human quantitative traits [20]. A model with two hidden layers and 64 units per layer generally displayed higher prediction accuracy than the others, namely a model consisting of a single hidden layer with 32 units per layer and a model consisting of five hidden layers with 32 units. In the Q200 scenario in our study, models with two hidden layers and more units per layer tended to exhibit higher accuracy (Figures 1–3), which is consistent with the human study.

It is unclear whether the increase in the numbers of hidden layers and units is proportional to the weights that are activated. It has been reported that a great part of weights included in huge neural networks such as convolutional neural networks can be pruned without affecting predictive performance [36,37]. A recent study noted that a benefit of large networks is increasing the chance to obtain a winning ticket (*i.e.*, a subset of networks that can achieve equivalent accuracy as the original model after training in isolation) from randomly initialized weights [38]. A core of the hypothesis is that the subnetwork is often unable to be trained in isolation from random initial values, thus necessitating appropriate initialization [38].

Figures S1 and S2 show the numbers and proportions of weights of which absolute values were more than 0.05 of the maximum absolute weights of each layer. The models were those with one and two hidden layers, and the applied scenarios were those with heritability = 1.0. Although the magnitudes differed among the models and scenarios considerably, redundancy in weights was observed as expected (Figure S1). In addition, two rough trends were observed: (1) as the number of units increased, the proportions of activated weights decreased (Figure S1), whereas the numbers increased (Figure S2); and (2) as the training population size grew, the proportions and numbers of activated weights decreased (Figures S1 and S2). The former occurred because the number of weights grew with greater magnitude than the decrease of the proportions of activated weights. As mentioned previously, an increase in the number of units was often accompanied by increased accuracy. Thus, the first trend suggests that the increased number of activated weights (or units) underpinned this increasing accuracy. However, it also can be deduced that most of the activated weights were redundant and that the redundant weights might remain activated (or stay around the initial values) because of shortage of the training population. Some of the activated weights, however, might include the winning tickets and result in higher accuracies. Greater training population sizes (probably much greater than 10,000) may be required to clarify the exact proportions of redundant weights.

The number of epochs that achieved the lowest MSEs in the validating populations, *i.e.*, the epoch number when optimized weights were used for prediction, displayed similar distributions among the training population sizes (Figure S3) and QTL-marker scenarios (Figure S4), whereas differences were observed among the models of neural networks (Figure S5). In each case, the epoch numbers accumulated in 10, which was the smallest number in our experiment setting (see Materials and Methods). This tendency was more frequently observed when small neural networks (*i.e.*, networks with a single hidden layer and/or 20 units per layer) were used (Figure S5). Figures S6–S8 show the scatter plots between the number of epochs and prediction accuracies. The smallest epoch number (10) was often related to extreme accuracies, *i.e.*, low (near 0) or high (>0.8) accuracies, and the frequency of the former case increased as the simulation scenarios became more difficult to predict, *i.e.*, as the training population size decreased, the numbers of markers and QTLs increased (Figures S6 and S7). The frequency also increased when small networks were applied (Figure S8). These results suggest that when small networks are used for complex data or data with a small training population size, learning tends to stop at extremely early steps. Two factors may be relevant to this issue, namely the learning rates of the Adam optimizer and the initialization methods for the weights. More careful tuning or choice may be required to solve this issue.

Dropout and one-hot genotype encoding were unable to increase the prediction accuracy (Figure S9). In ref. [20] the authors optimized the dropout rates using a genetic algorithm and found that the optimized rates were often 0.01, which was the lower bound of the optimization. Taken together with our results, it can be concluded that dropout would not be effective for genomic prediction. One-hot encoding was also reported by the authors [20]. Although the predictive performance of the model based on one-hot encoding depended on traits, the model occasionally exhibited the highest accuracy. The influence of genotype coding may need to be studied further.

### Comparison with random forests

The relative performance between neural networks and random forests depended on the simulation scenarios (Figures 1–3). In the Q20 scenario, neural networks exhibited higher accuracy when the heritability was 1.0 and displayed similar accuracy when the heritability was 0.2 or 0.6. In the Q20LM180 and Q20UM180 scenarios, when the heritability was 1.0, the prediction accuracies of neural networks depended on the training population size; specifically, neural networks generally exhibited higher, similar, and lower accuracies when the size was 10,000, 5000, and 1000, respectively, than random forests. When the heritability was 0.6 or 0.2 in the Q20LM180 and Q20UM180 scenarios, neural networks displayed comparative or inferior accuracies than random forests (Figures 2 and 3, respectively). In the Q200 scenario, neural networks and random forests generally exhibited similar accuracies. However, when the training population size was 10,000, neural networks with two hidden layers could provide slightly higher accuracies. These results suggest that neural networks are more sensitive to the training population size and heritability (the amount of noises) than random forests, although neural networks have the greater capability to predict epistatic genetic values.

**Figure 2.**
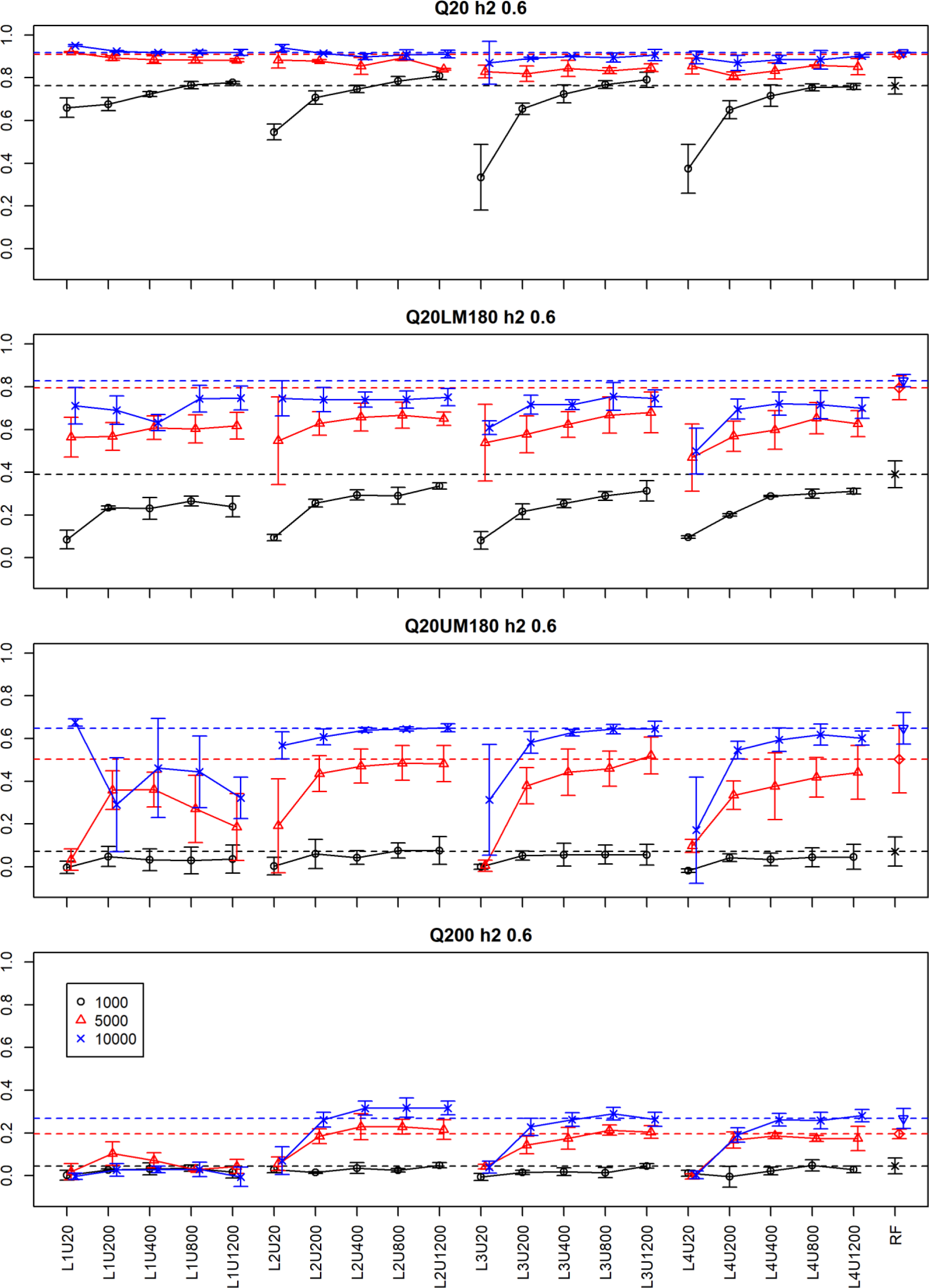
Prediction accuracies when the broad-sense heritability was 0.6. Four plots correspond with the four quantitative trait locus (QTL)-marker scenarios: Q20, Q20LM180, Q20UM180, and Q200 (see the text for the explanations of the QTL-marker scenarios). The y-axis is the prediction accuracy, and the x-axis presents the neural network models (L1U20 to L4U1200) and random forests (RFs). L and U in the model names represent the number of hidden layers and units per hidden layer, respectively. The dashed horizontal lines indicate the mean accuracies obtained using RFs. Colors and symbols indicate the training population sizes (1000, 5000, and 10,000). Bars are the standard deviations.

**Figure 3.**
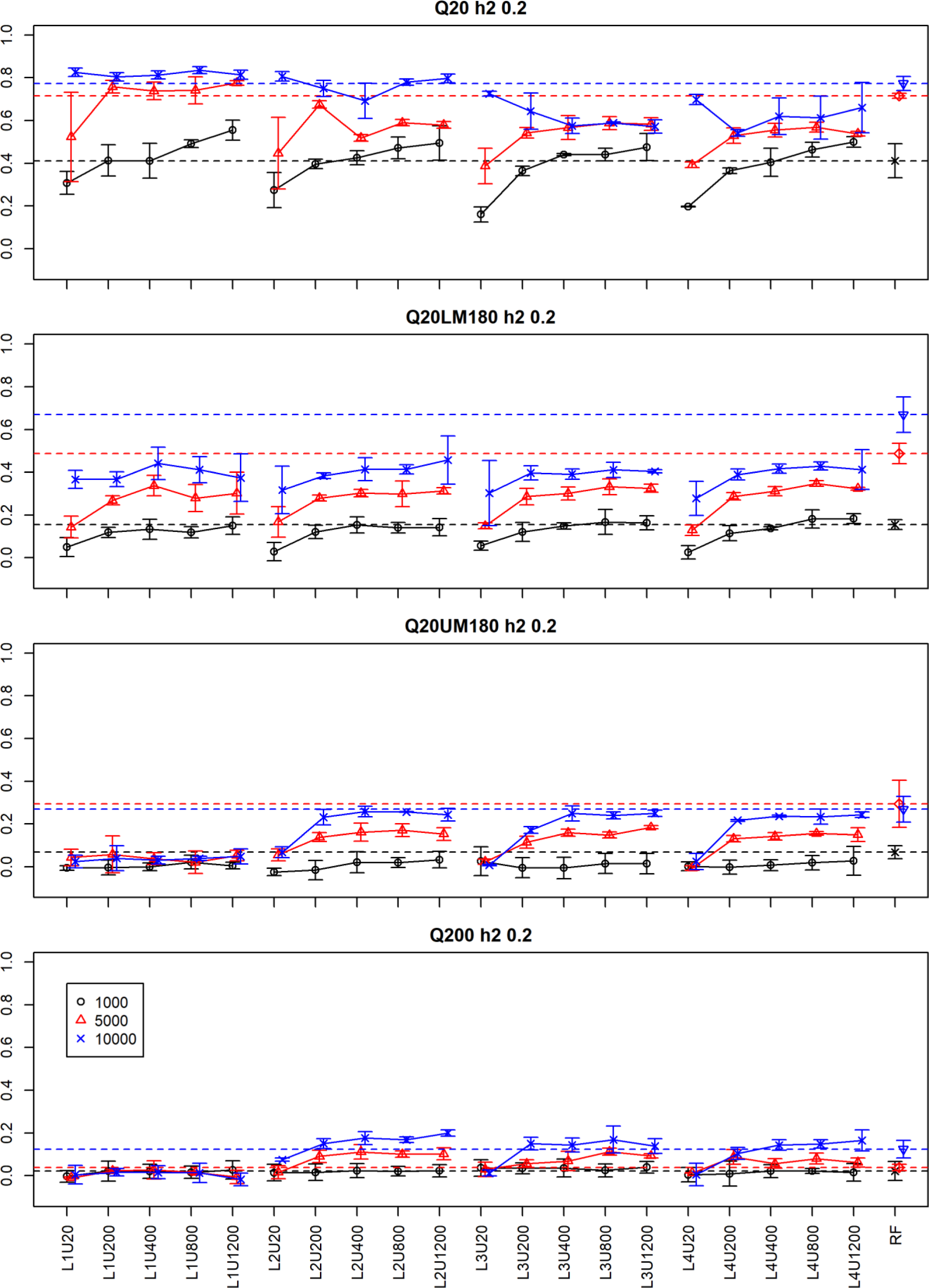
Prediction accuracies when the broad-sense heritability was 0.2. The four plots correspond with the four quantitative trait locus-marker scenarios: Q20, Q20LM180, Q20UM180, and Q200 (see the text for the explanations of the QTL-marker scenarios). The y-axis is the prediction accuracy, and the x-axis presents the neural network models (L1U20 to L4U1200) and random forests (RFs). L and U in the model names represent the number of hidden layers and units per hidden layer, respectively. The dashed horizontal lines indicate the mean accuracies obtained using RFs. Colors and symbols indicate the training population sizes (1000, 5000, and 10,000). Bars are the standard deviations.

The robustness of random forests to small training populations in genomic prediction was suggested in a previous study using simulations based on real rice data [39]. A comparison study between random forests and neural networks with a single hidden layer using multiple quantitative traits of crops showed that prediction accuracies were, on average, similar between the two methods [40]. Considering the small population sizes used in ref. [40] (at most 911), epistatic genetic values were probably unpredictable for the both methods, and prediction by these methods would have relied on additive and/or dominance components.

### Beyond genomic prediction

As described in the introduction, prediction of epistatic genetic values was examined in this study because interactions between explanatory variables are commonly observed in regression problems. In addition to epistasis, another example of interactions relating to breeding is the interaction between genes and environments, which is prominent particularly in plants. Whereas mixed effect models [41,42] and physiological model-based approaches [43,44] are often used to predict gene-by-environment interactions, several authors proposed neural networks that used both environmental covariates (*i.e.*, environment identifiers or climate covariates) and genotype information (*i.e.*., genotype identifiers, marker genotypes, or decomposed genetic relationship matrices) as inputs [45–47]. In ref. [45], relatively narrow (up to 100 units per layer) and shallow (up to three hidden layers) networks were applied to nine crop datasets without comparison with other non-linear methods. Given the sample sizes of the data (at most 2374) and the network structures, it might be infeasible for neural networks to exhibit better performance than random forests in terms of predicting gene-by-environment interactions. In another study, a different type of neural networks, namely long short-term memory-recurrent neural networks, was used to apply time series environment covariates for predicting soybean yields [46]. The properties of recurrent neural networks in predicting gene-by-environment interactions may differ from those of multilayer perceptron, and further studies will be needed for clarification. In ref. [47], a narrow and deep network (21 hidden layers and 50 units per layer) was used to predict the yield of maize. Although such a model structure is not supported from the results of the present study, the authors adopted the residual learning framework using shortcut connections, which realizes extremely deep architectures [48]. Considering the benefits of deeper architectures in complex problems [49], clarifying the effects of residual learning on the prediction of interactions between variables is worth further studies.

## Acknowledgments

The author thanks the office staff of the National Agriculture and Food Research Organization for their support on daily works.

## Data availability

The python and R scripts for the neural networks and random forests and the simulated datasets are available at https://github.com/Onogi/PredictingEpistaticGeneticValues.

## Notes

https://github.com/Onogi/PredictingEpistaticGeneticValues

