## Supplementary material for "Predicting genotypic values associated with gene interactions using neural networks: A simulation study for investigating factors affecting prediction accuracy": Figure S1

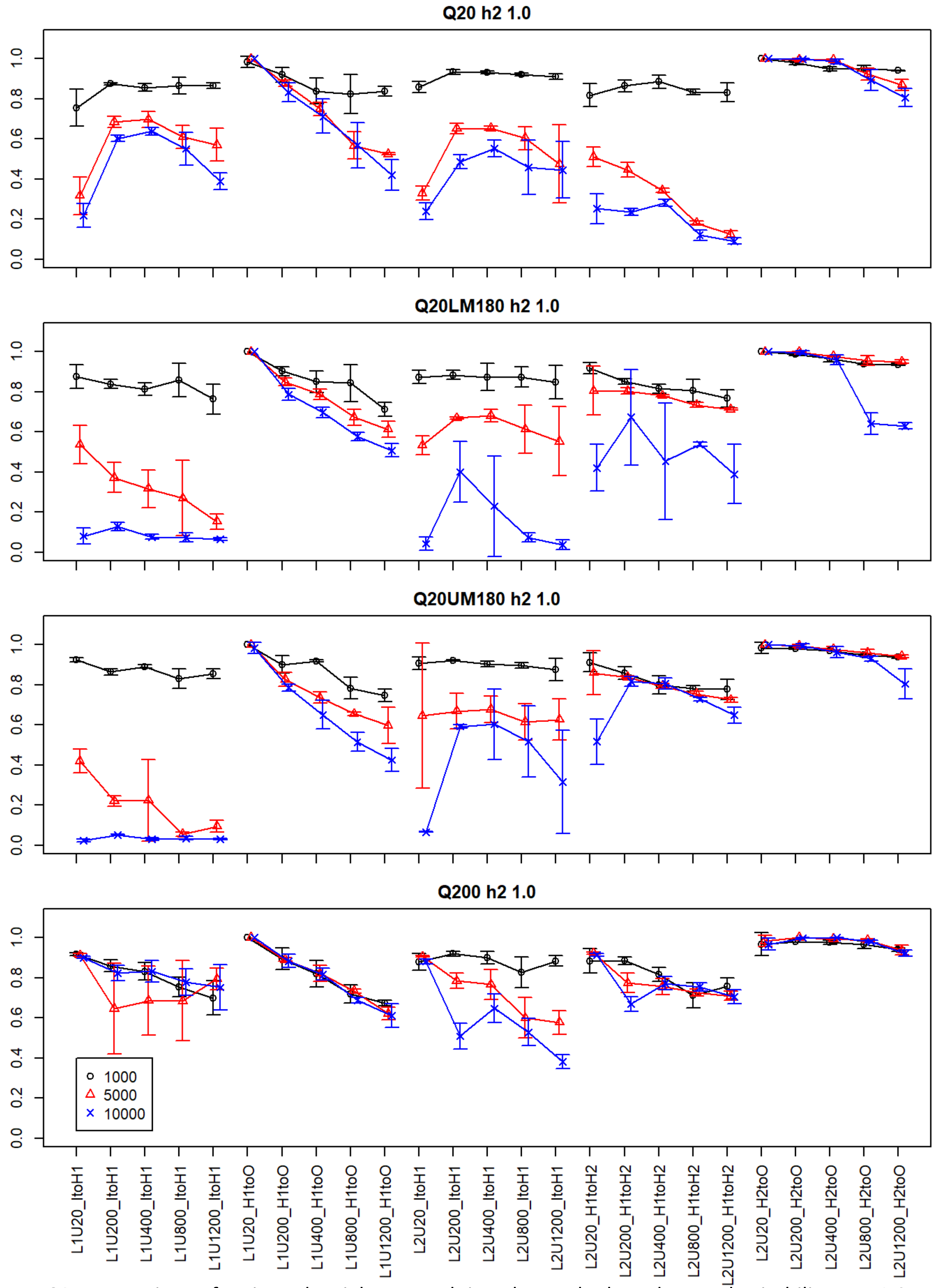


**Figure S1.** Proportions of activated weights at each interlayer. The broad-sense heritability was 1.0. Four plots correspond with four quantitative trait locus (QTL)-marker scenarios: Q20, Q20LM180, Q20UM180, and Q200 (see the text for the QTL-marker scenarios). The y-axis is the proportion, and the x-axis presents the neural network models (L1U20 to L2U1200) and interlayers. L and U in the model names represent the number of hidden layers and units per hidden layer, respectively. I, H1, H2, and O indicate the input, 1st and 2nd hidden layers, and output, respectively. Thus, ItoH1 indicates the weights connecting the input and 1st hidden layer. Colors and symbols indicate the training population sizes (1000, 5000, and 10,000). Bars are the standard deviations.
