## Supplementary material for "Predicting genotypic values associated with gene interactions using neural networks: A simulation study for investigating factors affecting prediction accuracy": Figure S3

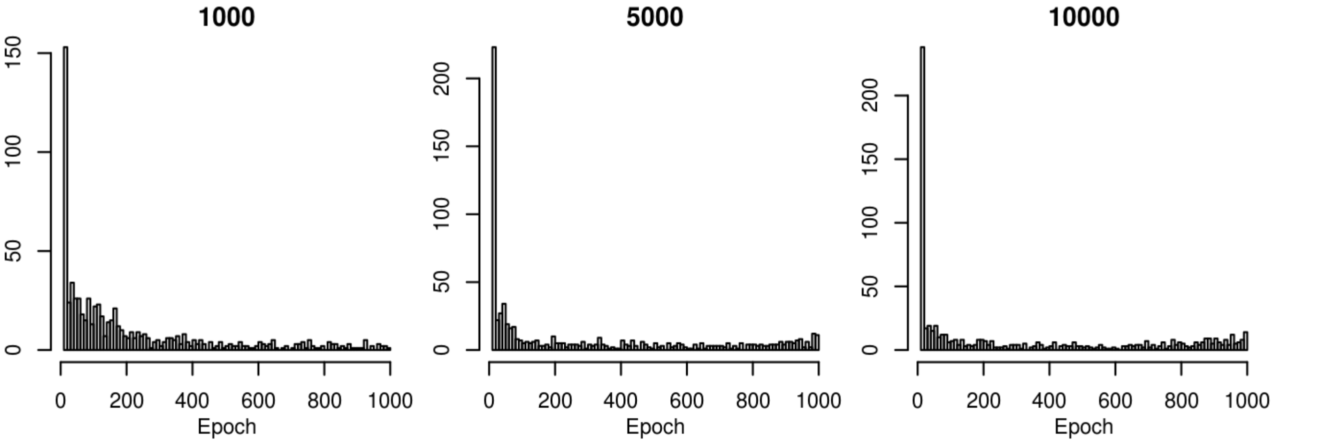


**Figure S3.** Epoch numbers of weights used for the prediction. The numbers were summarized for the training population size (1000, 5000, and 10,000). The y-axis is the frequency, and the x-axis is the epoch number.
