## Supplementary material for "Predicting genotypic values associated with gene interactions using neural networks: A simulation study for investigating factors affecting prediction accuracy": Figure S5

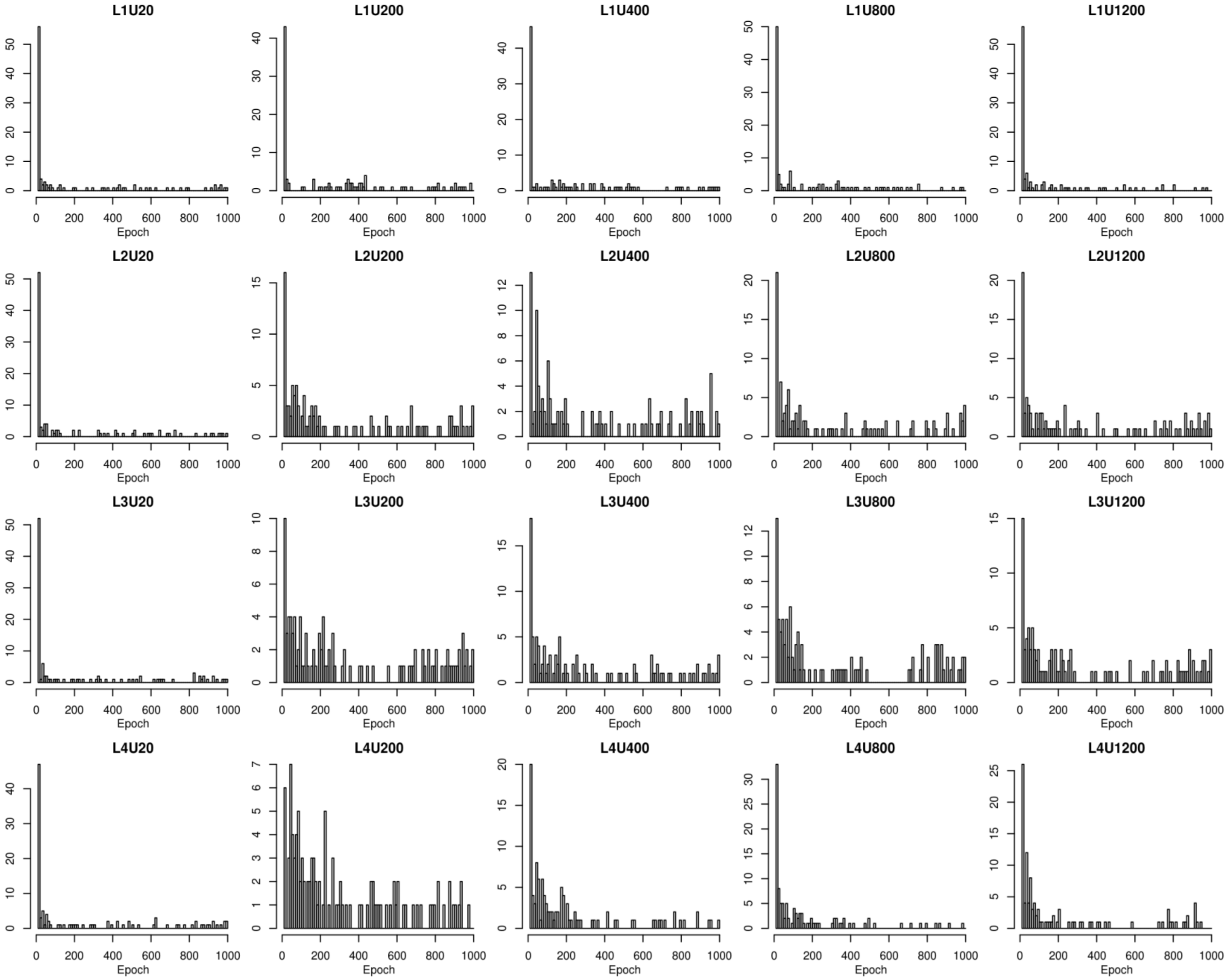


**Figure S5.** Epoch numbers of weights used for the prediction. The numbers were summarized for the neural network models. L and U represent the number of hidden layers and units per hidden layer, respectively. The y-axis is the frequency, and the x-axis is the epoch number.
