## Supplementary material for "Predicting genotypic values associated with gene interactions using neural networks: A simulation study for investigating factors affecting prediction accuracy": Figure S6

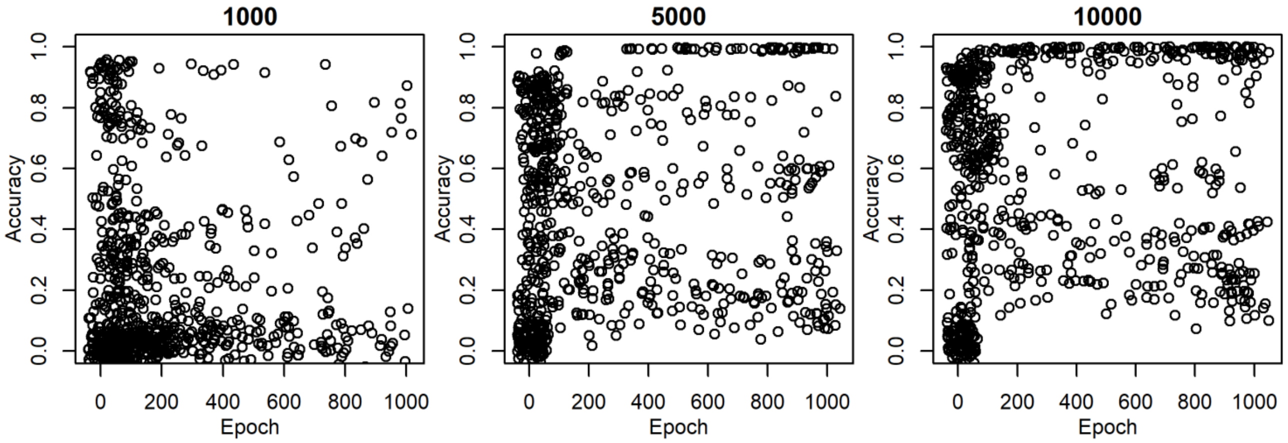


**Figure S6.** Scatter plots between the epoch numbers of weights used for the prediction and prediction accuracies. The numbers were summarized for the training population size (1000, 5000, and 10,000). The y-axis is the prediction accuracy, and the x-axis is the epoch number. The epoch numbers were jittered for visualization.
