## Supplementary material for "Predicting genotypic values associated with gene interactions using neural networks: A simulation study for investigating factors affecting prediction accuracy": Figure S7

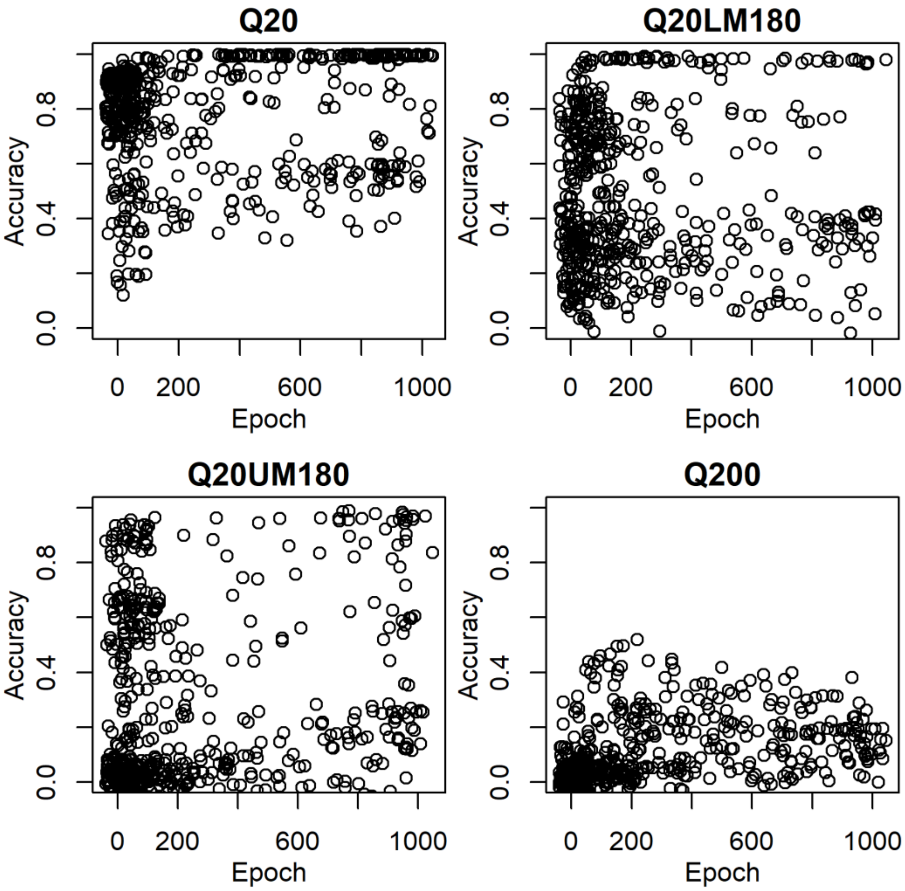


**Figure S7.** Scatter plots between the epoch numbers of weights used for the prediction and prediction accuracies. The numbers were summarized for the quantitative trait locus-marker scenarios (Q20, Q20LM180, Q20UM180, and Q200). The y-axis is the prediction accuracy, and the x-axis is the epoch number. The epoch numbers were jittered for visualization.
