## Supplementary material for "Predicting genotypic values associated with gene interactions using neural networks: A simulation study for investigating factors affecting prediction accuracy": Figure S9

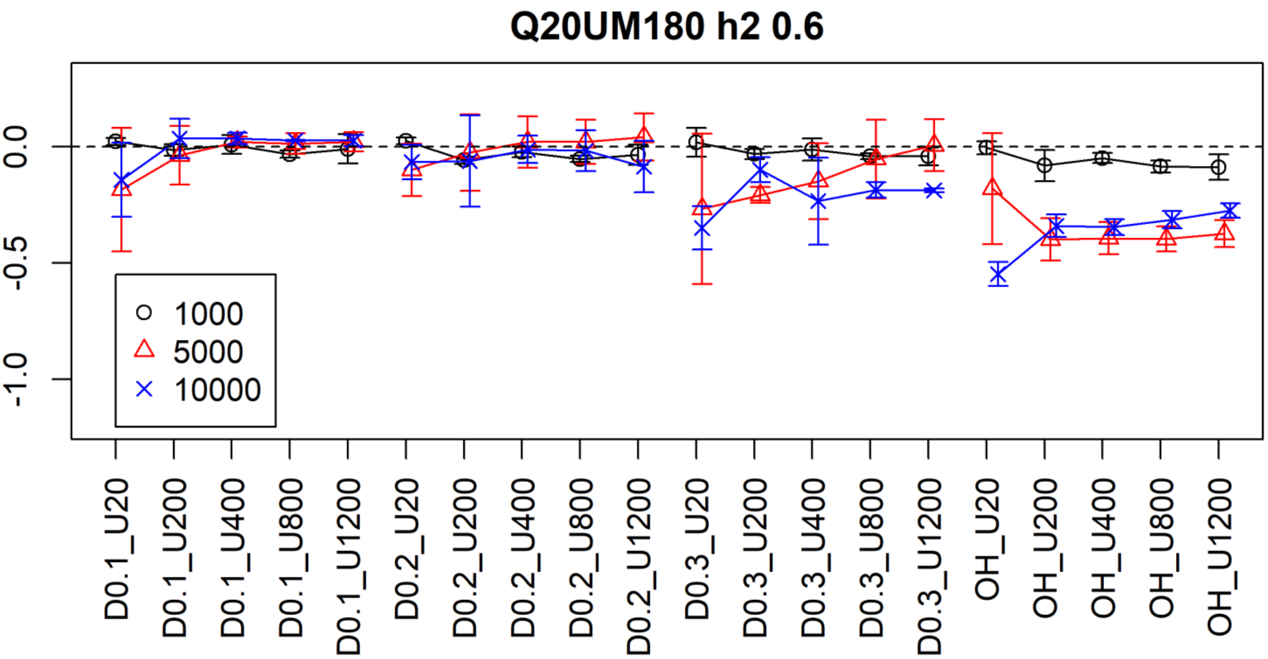


**Figure S9.** Differences in prediction accuracies between models with and without dropouts and between models with additive coding and one-hot encoding of genotypes. The y-axis presents the differences of prediction accuracies. Positive values indicate that the models with dropouts or models with one-hot encoding are better than those without dropout or those with additive coding. The x-axis presents the model names. D and U in the model names represent the dropout rates and the number of units per hidden layer, respectively. The number of hidden layers was two. OH indicates one-hot encoding. Colors and symbols indicate the training population sizes (1000, 5000, and 10,000). Bars represent the standard deviations.
